# Mitochondrial Function is Preserved Under Cysteine Starvation via Glutathione Catabolism in NSCLC

**DOI:** 10.1101/2022.10.06.511221

**Authors:** Nathan P. Ward, Sang Jun Yoon, Tyce Flynn, Amanda Sherwood, Juliana Madej, Gina M. DeNicola

## Abstract

Cysteine metabolism occurs across cellular compartments to support diverse biological functions and prevent the induction of ferroptosis. Though the disruption of cytosolic cysteine metabolism is implicated in this form of cell death, it is unknown whether the substantial cysteine metabolism resident within the mitochondria is similarly pertinent to ferroptosis. Here, we show that despite the rapid depletion of intracellular cysteine upon loss of extracellular cystine, cysteine-dependent synthesis of Fe-S clusters persists in the mitochondria of lung cancer cells. This promotes a retention of respiratory function and a maintenance of the mitochondrial redox state. Under these limiting conditions, we find that mitochondrial glutathione sustains the function of the Fe-S proteins critical to oxidative metabolism. This is achieved through CHAC1 catabolism of the cysteine-containing tripeptide within the mitochondrial matrix. We find that disrupting Fe-S cluster synthesis under cysteine restriction protects against the induction of ferroptosis, suggesting that the preservation of mitochondrial function is antagonistic to survival under starved conditions. Overall, our findings implicate mitochondrial cysteine metabolism in the induction of ferroptosis and reveal a novel mechanism of mitochondrial resilience in response to nutrient stress.

## Main

The proteinogenic amino acid cysteine (Cys) is a significant contributor to cellular functions due to its resident thiol group. Cysteine supplies this reactive sulfur in support of cellular antioxidant capacity (glutathione [GSH]), electron transfer (iron-sulfur [Fe-S] clusters), energy metabolism (coenzyme A), and osmoregulation (taurine)^1^. Despite the essentiality of cysteine for these processes, there is an apparent therapeutic window in restricting cysteine systemically as an anti-cancer strategy^2, 3^. Indeed, cysteine becomes conditionally essential across many tumor species^4^, in part due to an inability to synthesize the amino acid from methionine through the transsulfuration pathway^5^. This dictates a reliance on an extracellular source of cysteine, predominantly in its oxidized form of cystine (Cys_2_). Cysteine availability can be targeted through its systemic depletion^2^ or inhibition of the primary Cys_2_ transporter, system X_c_^-6^.

Cysteine restriction promotes an iron-dependent form of cell death known as ferroptosis^7^, where Fenton chemistry promotes the generation of free radicals that initiate lipid radical formation and the unsustainable peroxidation of plasma membrane phospholipids^8^. The resulting loss of membrane integrity is suppressed by intrinsic pathways mediated by ferroptosis suppressor protein 1 (FSP1)^9^ and glutathione peroxidase 4 (GPX4)^10^ to ward off ferroptosis under basal conditions. However, in the absence of cysteine, cancer cells are unable to synthesize GSH, restricting GPX4 activity and inducing ferroptosis.

Additional factors have been implicated in the induction of ferroptosis^11^, including the engagement of mitochondrial metabolism and the tricarboxylic acid (TCA) cycle^12, 13^. Mechanisms by which the TCA cycle promotes ferroptosis include the generation and export of citrate to drive the synthesis of new substrates for lipid peroxidation^14^, and the stimulation of the electron transport chain (ETC)^13^. However, how ETC activity promotes the induction of ferroptosis remains to be fully elucidated^15^. Existing evidence suggests that ATP production^14^ and the generation and extramitochondrial release of reactive oxygen species (ROS) may be relevant consequences^15, 16^.

One aspect of mitochondrial metabolism that has not been adequately addressed is the role of resident cysteine metabolism within the mitochondria. Though the disruption of cytosolic cysteine metabolism (i.e., GSH synthesis) plays a significant role in the induction of ferroptosis under cysteine limitation^5, 17^, it remains unknown if alterations in mitochondrial cysteine metabolism are also operative. Mitochondrial cysteine is used in the synthesis of mitochondrially encoded protein components of the ETC, hydrogen sulfide production^18^, and Fe-S cluster synthesis^19^ in support of bioenergetic metabolism. Given the established connection between oxidative metabolism and ferroptosis^12, 13^, the requirement for cysteine in support of respiration suggests it may be particularly relevant to ferroptosis. Furthermore, a disruption in mitochondrial cysteine metabolism could explain the gradual loss of mitochondrial function reportedly associated with prolonged cysteine deprivation^3, 13, 20, 21^.

To interrogate the influence of mitochondrial cysteine metabolism in the induction of ferroptosis, we evaluated mitochondrial respiratory function in response to prolonged cysteine deprivation with regard to the capacity for Fe-S cluster synthesis. We evaluated this in the context of non-small cell lung cancer (NSCLC), given their sensitivity to cysteine starvation^22^ and the robust mitochondrial metabolism exhibited by human lung tumors^23, 24^. Remarkably, despite the rapid depletion of mitochondrial cysteine under starved conditions, Fe-S cluster synthesis persisted in support of respiratory function through the onset of ferroptosis. This was achieved through the catabolism of mitochondrial GSH and at the expense of NSCLC viability.

## Results

### Mitochondrial Respiratory Function Persists Under Cystine Deprivation

To determine the influence of cysteine deprivation on mitochondrial metabolism, we restricted extracellular cystine in a panel of NSCLC cells. We opted to starve cells of their source of cysteine rather than inhibit cystine uptake with erastin to directly assess the consequences of cysteine deprivation without the confounding influence of erastin’s promiscuous inhibition of the mitochondrial voltage-dependent anion channel^25, 26^. As previously described, NSCLC cells succumb to ferroptosis in the absence of cysteine^5, 27^ (**Extended Data Figure 1a-c**). We begin to observe the accumulation of ferroptotic cells, as detected upon the intercalation of a cell-impermeable fluorescent dye with nuclear DNA, following 24h of cystine deprivation (**Extended Data Figure 1a**). Therefore, we performed our mitochondrial analyses upon 20h of starvation to ensure sufficient nutrient stress in the absence of a loss of cell viability.

To assess general mitochondrial respiratory function under cystine deprivation, we employed the Seahorse MitoStress test. Generally, the basal oxygen consumption rate (OCR) of NSCLC cells was not diminished in the absence of cystine (**Figure 1a**), indicating a retention of mitochondrial respiration. This coincided with a maintenance of mitochondrial coupling efficiency (**Figure 1b**), an indicator of the functionality of the mitochondrial inner membrane (IMM)^28^. Further, though it was reported that the IMM becomes dramatically hyperpolarized under cysteine deprivation^13^, mitochondrial membrane potential (ΔΨ_m_) was either minimally increased or unchanged under starved conditions (**Figure 1c**). Consistent with the preservation of a functional IMM, ATP-linked respiration was not compromised in the absence of cystine (**Figure 1d**). Moreover, several NSCLC lines exhibited a significant elevation in maximal respiratory capacity that could not be explained by an increase in mitochondrial density (**Figures 1e and 1f**), suggesting that cysteine deprivation may enhance mitochondrial function beyond the requirement for oxidative phosphorylation. Collectively, these data indicate a robust persistence of mitochondrial respiration under cystine deprivation that occurs independent of an expected insult to the IMM^13, 20^.

**Figure 1:**
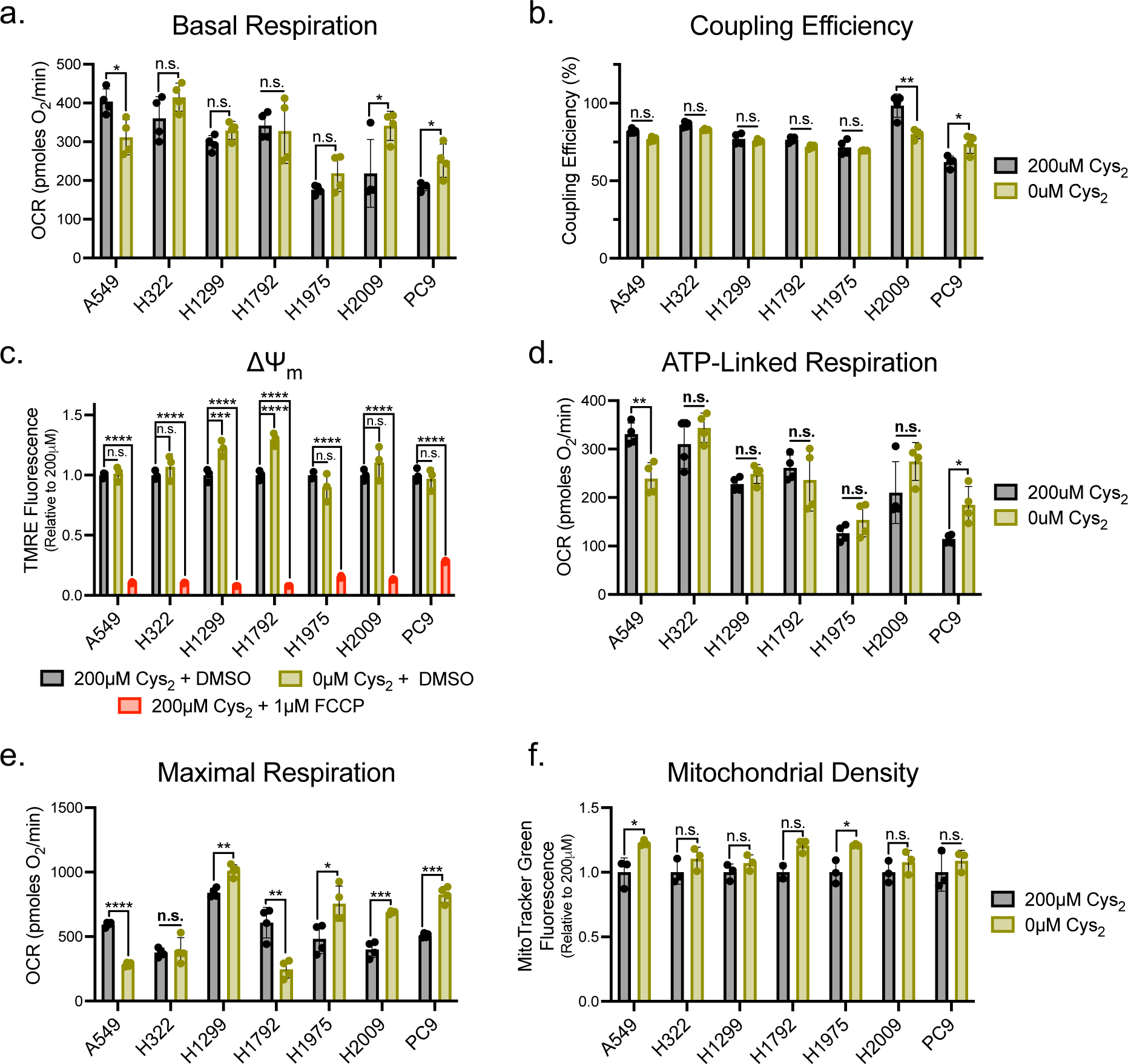
Mitochondrial Respiration is Sustained in the Absence of Extracellular Cys_2_. **a**, Basal oxygen consumption rate of NSCLC cells stimulated with 10mM glucose and 2mM L-glutamine following culture in the presence or absence of cystine (n=4 per condition). **b**, Percentage of basal oxygen consumption linked to the generation of ATP in NSCLC cells fed or starved of cystine (n=4 per condition). **c**, Average relative TMRE fluorescence of NSCLC cells cultured with indicated [cystine] or treated with FCCP for 15 minutes (n=3 per condition). **d**, Oligomycin-sensitive oxygen consumption rate of cystine-fed or starved NSCLC cells (n=4 per condition). **e**, FCCP-stimulated oxygen consumption rate in in NSCLC cells fed or starved of cystine (n=4 per condition). **f**, Average relative MitoTracker Green fluorescence of NSCLC cells cultured in the presence or absence of cystine (n=3 per condition). For all experiments, cells were cultured in the indicated [cystine] for 20 hours prior to the indicated analysis. Data represent mean values ± s.d; n.s., not significant, *, P<0.05, **, P<0.01, ***, P<0.001, ****, P<0.0001. For **a**, **b**, **d-f**, P values were calculated using unpaired Student’s *t*-test. For **c**, P values were calculated using a one-way ANOVA. All data are representative of at least 3 experimental replicates.

### Cystine Deprivation Does Not Promote Mitochondrial Oxidative Stress

To confirm that the persistence of respiratory function is not damaging to the mitochondria of NSCLC cells, we interrogated the mitochondrial redox state in response to cystine starvation. First, we employed a series of fluorescent ROS indicators to assess changes in the mediators of oxidative stress. While we observe the hallmark increase in lipid peroxidation at the cellular level (**Figure 2a**)^8^, and a coincident increase in cellular free radical levels under cystine deprivation (**Figure 2b**), these indications of oxidative stress do not extend to the mitochondria. Use of specific and mitochondrially targeted indicators of superoxide (•O_2_^-^) and hydrogen peroxide (H_2_O_2_) revealed that prolonged cystine starvation did not alter the abundance of either species (**Figures 2c and 2d**). This suggests that cysteine deprived mitochondria can scavenge the significant ROS generated as a byproduct of persistent mitochondrial metabolism^29^.

**Figure 2:**
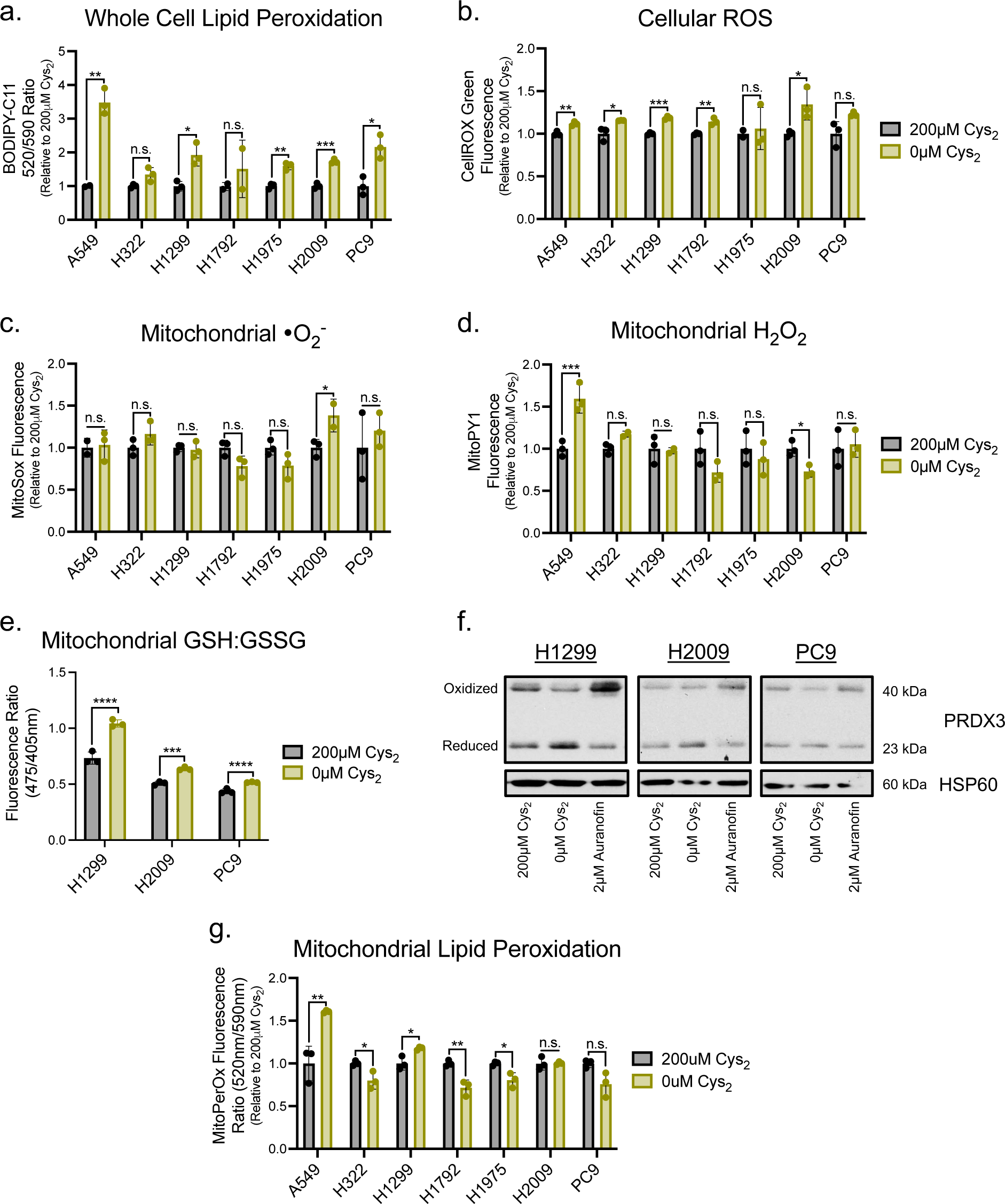
Cys_2_ Starvation Does Not Promote Mitochondrial Oxidative Stress. **a-d**, Following culture in the presence or absence of cystine, NSCLC cells were treated with ROS-sensitive fluorescent dyes to determine the average relative levels of **a**, cellular membrane lipid peroxides, **b**, cellular ROS, **c**, mitochondrial superoxide, or **d**, mitochondrial hydrogen peroxide (n=3 per condition for each experiment). **e**, Ratio of Mito-Grx1-roGFP2 fluorescence as indicative of the proportion of the biosensor bound to reduced (GSH) or oxidized (GSSG) glutathione in H1299, H2009, and PC9 cells cultured with or without cystine for 16 hours (n=3 per condition). **f**, Redox immunoblotting of HSP60 and the PRDX3 oxidation state in H1299, H2009, and PC9 cells following culture with the indicated [cystine] or 1 hour treatment with auranofin. **g**, Relative MitoPerOX fluorescence ratio of NSCLC cells cultured in the presence or absence of cystine to indicate the extent of mitochondrial membrane lipid peroxidation (n=3 per condition). For **a-d**, **f**, and **g** cells were cultured in the indicated [cystine] for 20 hours prior to the indicated analysis. Data represent mean values ± s.d; n.s., not significant, *, P<0.05, **, P<0.01, ***, P<0.001, ****, P<0.0001. For **a**-**e**, and **g**, P values were calculated using two-tailed unpaired Student’s *t*-test. All data are representative of at least 3 experimental replicates.

Mitochondrial ROS metabolism is achieved predominantly through the activity of two complementary, yet distinct thiol-based antioxidant systems^30^. Both the GSH tripeptide and the small molecular weight protein thioredoxin (TXN) support ROS detoxification by a network of antioxidant proteins within the mitochondria^31^. Through use of a genetically encoded and mitochondrially targeted biosensor^32^, we determined that the mitochondrial GSH pool was more reduced under cystine deprivation relative to replete conditions (**Figure 2e**). Further, an assessment of the oxidation state of peroxiredoxin 3 (PRDX3), a component of the TXN system, revealed an increase in the functional reduced form upon cystine starvation (**Figure 2f**). Together these data suggest that cystine deprivation elicits a more reduced mitochondrial matrix despite robust metabolic activity. In agreement with this apparent lack of oxidative stress, we did not observe mitochondrial lipid peroxidation upon cystine deprivation (**Figure 2g**), which indicates the increase in whole cell lipid peroxidation is confined to the plasma membrane (**Figure 2a**). Collectively, these data reflect that a maintenance of mitochondrial redox homeostasis accompanies the persistence of respiration in cysteine restricted NSCLC cells.

### Fe-S Cluster Synthesis is Sustained Under Cystine Deprivation

Cysteine is a requirement for sustained mitochondrial function, due in large part to its utilization in the synthesis of Fe-S clusters. These redox cofactors mediate electron transfer and support the enzymatic function of proteins critical to mitochondrial metabolism, including components of the ETC and TCA cycle^19^. Fe-S cluster synthesis is initiated in the mitochondria and requires the coordination of iron and cysteine-derived sulfur by a multi-protein complex (**Figure 3a**). Disrupting this machinery is associated with the loss of ETC function and an increase in ROS production^33, 34^. Furthermore, deficiencies in Fe-S cluster synthesis are associated with severe mitochondrial defects underlying varied human pathologies^35^. Considering this, the absence of mitochondrial dysfunction in NSCLC cells under cystine starvation suggests that Fe-S cluster synthesis is sustained.

**Figure 3:**
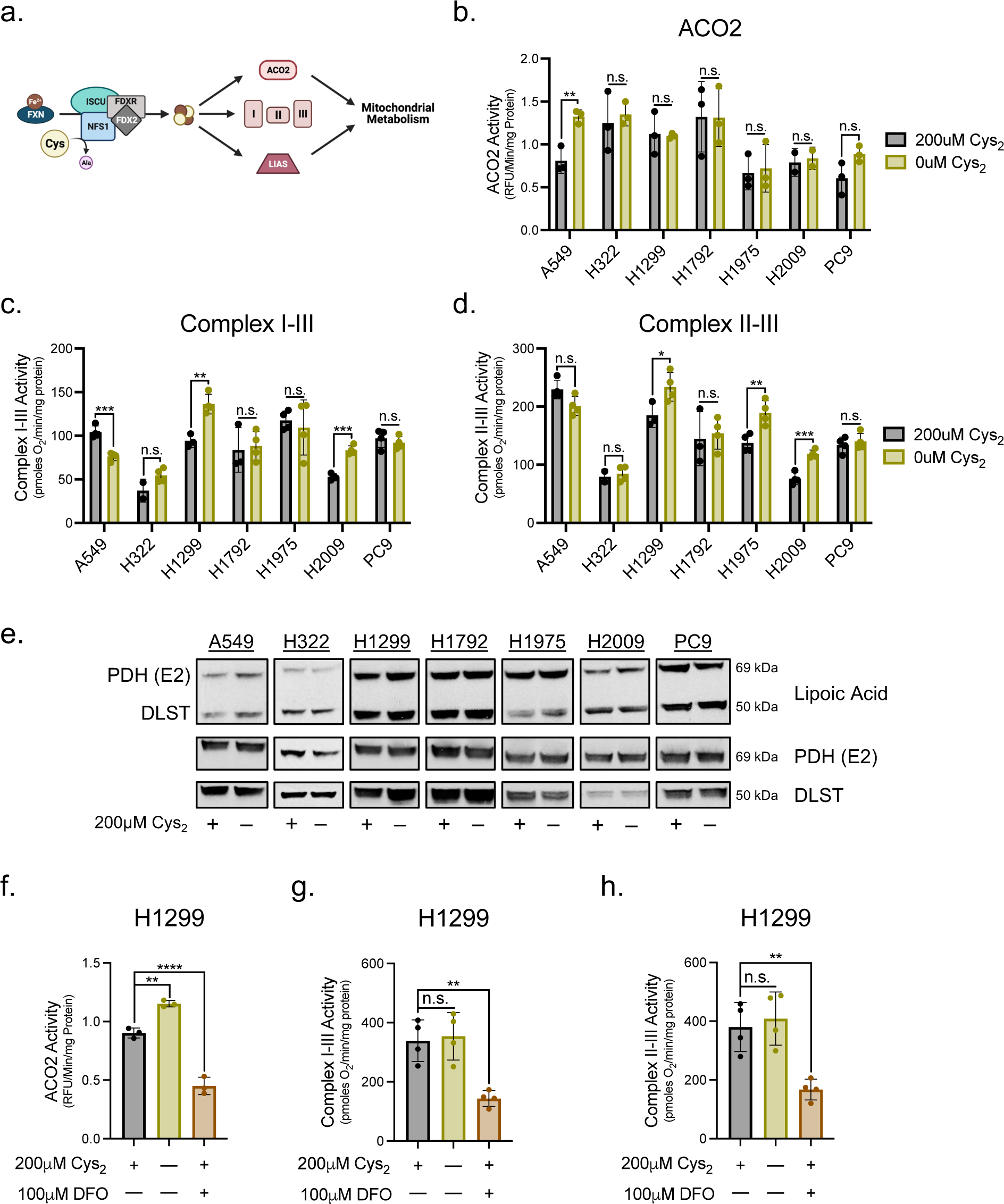
Fe-S Protein Function is Preserved Under Cys_2_ Deprivation. **a**, Schematic of Fe-S cluster synthesis and its contribution to enzymatic function that supports mitochondrial metabolism. **b**, ACO2 activity in mitochondrial lysates of NSCLC cells fed or starved of cystine (n=3 per condition). **c**, Supercomplex I-III activity in permeabilized NSCLC cells stimulated with 10mM pyruvate and 1mM malate following culture in the presence or absence of cystine (n :=3 per condition). **d**, Supercomplex II-III activity in permeabilized NSCLC cells subject to rotenone inhibition of complex I and stimulated with 10mM succinate following culture in the presence or absence of cystine (n :=3 per condition). **e**, Immunoblot analysis of PDH complex subunit E2 and α-ketoglutarate dehydrogenase subunit DLST lipoylation following culture in cystine replete or starved conditions. **f**, ACO2 activity in H1299 cells fed/starved of cystine or treated with the iron chelator, DFO (n=3 per condition). **g** and **h**, ETC supercomplex activities in permeabilized H1299 cells following culture in the presence or absence of cystine or treatment with DFO. For all experiments, cells were cultured in treatment conditions for 20 hours prior to the indicated analysis. Data represent mean values ± s.d; n.s., not significant, *, P<0.05, **, P<0.01, ***, P<0.001, ****, P<0.0001. For **b-d**, P values were calculated using two-tailed unpaired Student’s *t*-test. For **f-h**, P values were calculated using a one-way ANOVA. All data are representative of at least 3 experimental replicates.

To scrutinize this possibility, we evaluated the functionality of mitochondrial Fe-S proteins under cystine deprivation. First, an assessment of the TCA cycle enzyme aconitase (ACO2) in isolated mitochondria showed no change in ACO2 activity upon cystine starvation (**Figure 3b**). Next, we employed a specialized Seahorse-based assay to specifically assess the function of the Fe-S cluster dependent complexes of the ETC^36, 37^. This revealed that the activities of the respiratory supercomplexes I-III and II-III were not diminished in the absence of extracellular cystine (**Figures 3c and 3d**). Lastly, we interrogated the function of the unique Fe-S protein lipoic acid synthetase (LIAS). Rather than simply mediate its catalysis, LIAS’ resident Fe-S cluster is consumed during its synthesis of lipoate moieties that facilitate metabolic protein function^38^. We evaluated the lipoylation status of the E2 component of the pyruvate dehydrogenase complex (PDH-E2) and dihydrolipoamide S-succinyltransferase (DLST) as surrogate markers for LIAS activity and found no change in protein lipoylation with respect to cysteine availability (**Figure 3e**). Collectively these data demonstrate a complete maintenance of Fe-S protein function under cystine starvation, suggesting that Fe-S cluster synthesis is sustained despite the absence of cysteine.

Still, there remained the possibility that Fe-S cluster turnover exceeds the 20h starvation timeframe used for these studies, which would negate the necessity of *de novo* synthesis. To interrogate this, we chelated cellular iron with deferoxamine (DFO) to manipulate the other input of Fe-S cluster synthesis^38^. In contrast to cystine starvation, 20h of iron chelation significantly diminished Fe-S protein function in H1299 cells (**Figures 3f-h**). Importantly, DFO treatment promoted a progressive decrease in respiratory complex function (**Extended Data Figures 2a and 2b**). This suggests that rather than stripping iron from existing clusters and rendering them inactive, iron chelation restricts synthesis of new clusters. Together, these data indicate that NSCLC cells retain the capacity to synthesize Fe-S clusters in the absence of an extracellular cysteine source.

### CHAC1 Supports Fe-S Protein Function Under Cystine Deprivation

The maintenance of Fe-S cluster synthesis under cystine starvation argues for the existence of an alternative source of mitochondrial cysteine. Given the absence of mitochondrial oxidative stress and more reduced state of the matrix under cystine starvation (**Figure 2**), we considered the possibility that the antioxidant function of GSH was dispensable, and that the tripeptide could serve as a mitochondrial cysteine sink. Intriguingly, characterization of the transcriptional response to cysteine deprivation has revealed glutathione-specific gamma-glutamylcyclotransferase 1 (CHAC1) as the most highly induced gene^6^. CHAC1 is a component of the intracellular GSH cleavage system (**Figure 4a**), which catabolizes GSH to yield 5-oxoproline and cysteinylglycine (Cys-Gly)^39^. Cys-Gly can then be further cleaved by various peptidases to release its constituent amino acids^40^. Considering this, we hypothesized that CHAC1 induction under cystine starvation would promote cysteine mobilization from GSH to support Fe-S cluster synthesis.

**Figure 4:**
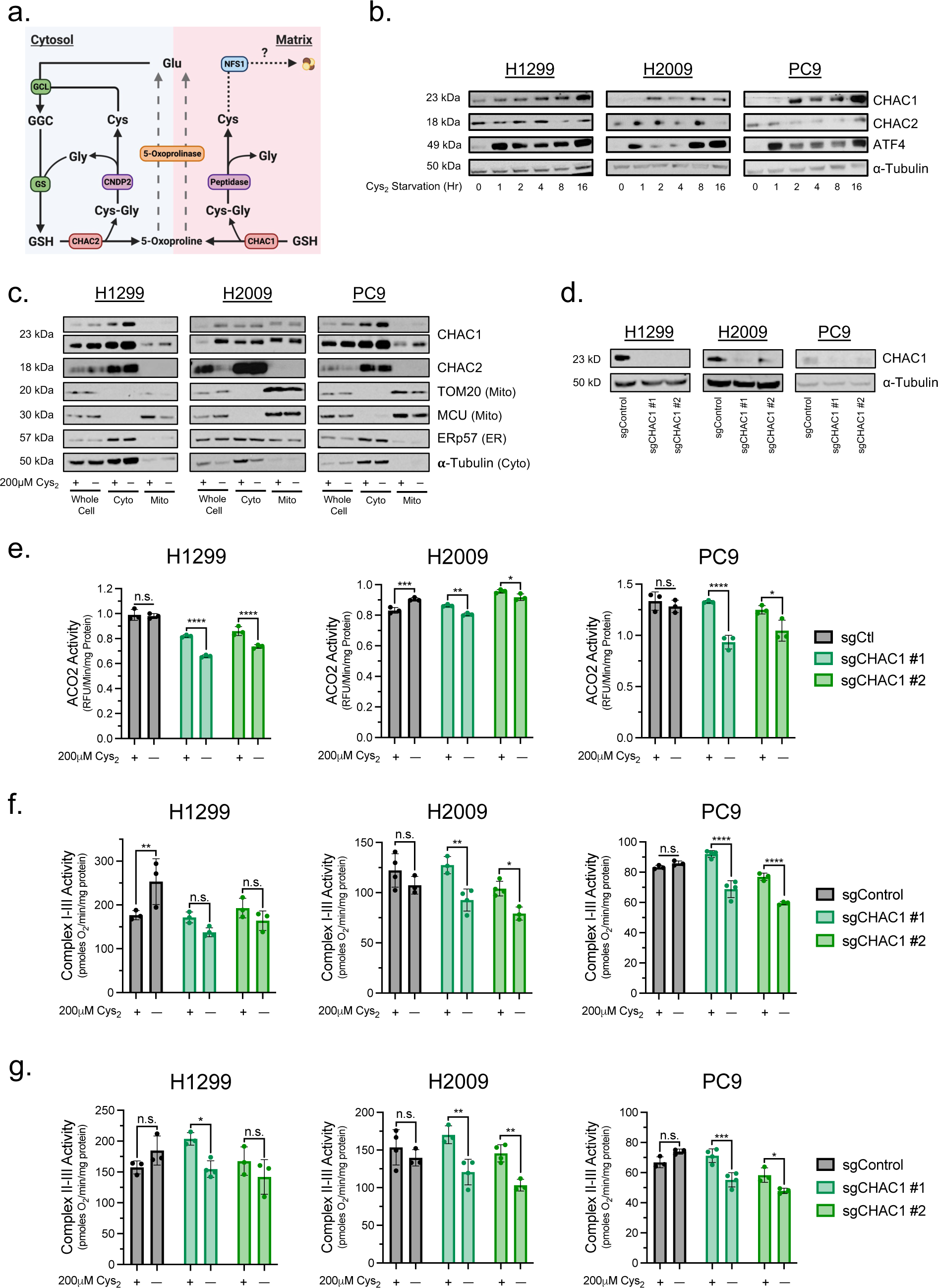
CHAC1 Loss Diminishes Fe-S Protein Function Under Cys2 Starvation. **a**, Schematic of the intracellular GSH cleavage system. **b**, Time-course immunoblot analysis of CHAC1, CHAC2, ATF4, and α-Tubulin expression in H1299, H2009, and PC9 cells under cystine starvation. **c**, immunoblot analysis of CHAC1, CHAC2, ERp57, MCU, TOM20, and α-Tubulin expression in either whole cell lysates or in fractionated lysates upon mitochondrial immunoprecipitation from H1299, H2009, and PC9-HA-Mito cells cultured in the presence or absence of cystine. **d**, immunoblot analysis of CHAC1 and α-Tubulin expression in H1299, H2009, and PC9 cells subject to CRISPR/Cas9-mediated knockout of CHAC1. **e**, ACO2 activities in CHAC1-deficient H1299, H2009, and PC9 cells subject to either cystine replete or starved conditions (n=3 per condition). **f** and **g**, ETC supercomplex activities in permeabilized CHAC1-deficient cells stimulated with their associated substrates following culture in the presence of absence of cystine (n :=3 per condition). For **c**, **e-g**, cells were cultured in the indicated [cystine] for 20 hours prior to analysis. Data represent mean values ± s.d; n.s., not significant, *, P<0.05, **, P<0.01, ***, P<0.001, ****, P<0.0001. For **e-g**, data are representative of at least 3 experimental replicates and P values were calculated using a two-way ANOVA.

In our NSCLC cells, we find that cystine starvation elicited a temporal accumulation of CHAC1 protein that coincides with an induction of ATF4 (**Figure 4b**), agreeing with previous reports that CHAC1 is an ATF4 target^39, 41^. In contrast, expression of CHAC2, the other intracellular mediator of GSH catabolism was not induced by cystine starvation (**Figure 4b**). Several expansive studies mapping the subcellular distribution of the human proteome have suggested that CHAC1 localizes to multiple organelles, including the mitochondria^42, 43^. We generated cells expressing a His-tagged construct within the outer mitochondrial membrane (HA-Mito)^44^ to facilitate the purification of mitochondria from cell homogenates and assess CHAC1 expression within the organelle. We found that CHAC1 is expressed within the mitochondria of NSCLC cells and that CHAC1 expression within the mitochondria was also increased under cystine starvation (**Figure 4c**). Importantly, CHAC2 does not localize to the mitochondria irrespective of cystine availability, suggesting any effect of GSH catabolism on mitochondrial function is attributable to CHAC1.

With this confirmation of mitochondrial localization, we next employed the CRISPR/Cas9 system^45^ to generate CHAC1 knockout cell lines to determine if CHAC1 influences Fe-S protein function by influencing cysteine availability (**Figure 4d**). We found that CHAC1 knockout had no detrimental effect on Fe-S protein function in the presence of cystine (**Figures 4e-f**). However, unlike in control cells, the activities of ACO2 and the Fe-S dependent respiratory supercomplexes were diminished to varying degrees under cystine starvation in CHAC1 knockout cells (**Figures 4e-f**). To supplement these findings, we also assessed the influence of CHAC1 expression on Fe-S protein function with a doxycycline-inducible shRNA system (**Extended Data Figure 3a**)^46^. Again, loss of CHAC1 protein did not influence ACO2 or ETC complex activity in the presence of cystine (**Extended Data Figures 3b-d**). Yet, we did observe a significant reduction in the functionality of these Fe-S proteins under cystine deprivation upon CHAC1 knockdown (**Extended Data Figures 3b-d**). To ensure that these effects were specific to CHAC1, we also evaluated the influence of CHAC2 knockdown on Fe-S protein function (**Extended Figure 3e**). Critically, CHAC2 loss had no effect on ACO2 or respiratory chain activity regardless of cystine availability (**Extended Data Figures 3f-h**). Collectively these data indicate that CHAC1 expression is enhanced in the mitochondrial matrix of cystine starved NSCLC cells in a manner that supports the function of Fe-S proteins. Further, this activity is confined to CHAC1 as its homolog CHAC2 is absent from the mitochondria.

### Mitochondrial GSH Supports Fe-S Protein Function in the Absence of Extracellular Cystine

To determine if CHAC1 has a discernable influence on the pools of mitochondrial cysteine and GSH in the potential support of Fe-S cluster synthesis, we endeavored to monitor the availability of these metabolites in response to cystine starvation (**Figure 5a**). To accomplish this, we employed an established liquid chromatography-mass spectroscopy (LC-MS) metabolomics scheme that makes use of HA-Mito cells to facilitate the rapid isolation of mitochondria for analysis of matrix metabolite^44, 47^. Though this system enables the detection of the wide range of metabolites present within the mitochondria, the nature of this scheme prevents the detection of cysteine due to the high reactivity of its resident thiol^44, 48^. To overcome this technical limitation, we utilized the alkylating agent n-ethylmaleimide (NEM)^49^ to stabilize thiol containing metabolites. Application of NEM rapidly alkylated cellular thiols, indicated by the stabilization of reduced PRDX3 within 30 seconds of treatment (**Extended Data Figure 4a**). Importantly, this did not interfere with the capacity to isolate mitochondria from HA-Mito cells (**Extended Data Figure 4b**).

**Figure 5:**
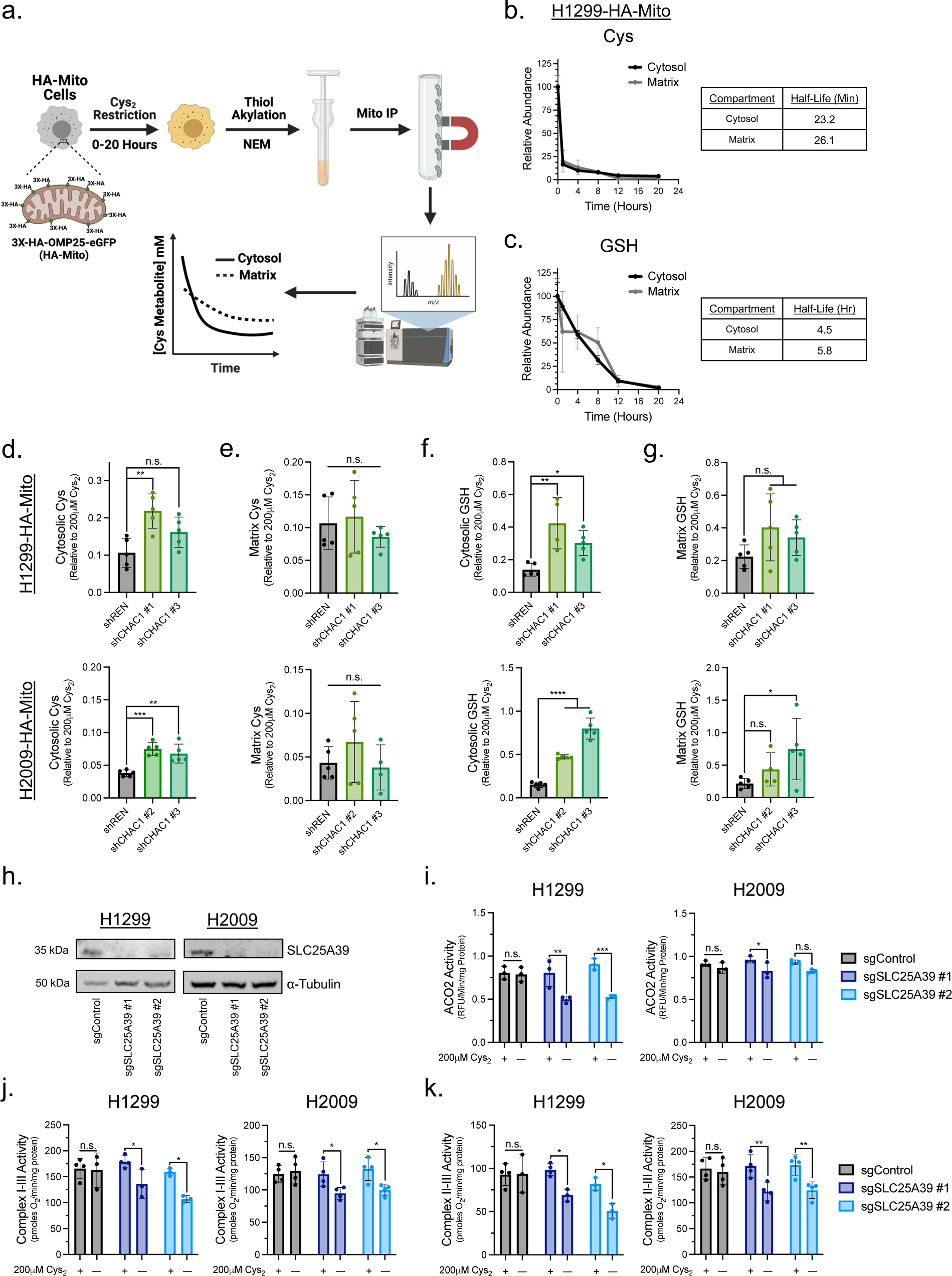
CHAC1 Promotes Cellular GSH Depletion Under Cys_2_ Deprivation. **a**, Schematic workflow of mitochondrial isolation coupled to LC-MS for the compartmentalized detection of thiol-containing metabolites. **b**, Determination of the half-life of cytosolic and matrix Cys in H1299-HA-Mito cells cultured in the absence of extracellular Cys for 20h (n=3 biological replicates per time point). **c**, Determination of the half-life of cytosolic and matrix GSH in H1299-HA-Mito cells cultured in the absence of extracellular Cys for 20h (n=3 biological replicates per time point). **d-f**, H1299 and H2009-HA-Mito cells were treated with 0.2μg/mL doxycycline for 5 days to induce shRNA knockdown of CHAC1 and then subject to cystine starvation for 12 hours. The pool sizes of **d**, cytosolic Cys, **e**, matrix Cys, **f**, cytosolic GSH, and **g**, matrix GSH were then determined relative to cystine replete conditions (n=5 biological replicates per condition). **h**, immunoblot analysis of SLC25A39 and α-Tubulin expression in H1299 and H2009 cells subject to CRISPR/Cas9-mediated knockout of SLC25A39. **i**, ACO2 activities in SLC25A39-deficient H1299 and H2009 cells subject to either cystine replete or starved conditions (n=3 per condition). **j** and **k**, ETC supercomplex activities in permeabilized SLC25A39-deficient cells stimulated with **j**, 10mM pyruvate and 1mM malate or **k**, 10mM succinate following culture in the presence of absence of cystine (n :=3 per condition). Data represent mean values ± s.d; n.s., not significant, *, P<0.05, **, P<0.01, ***, P<0.001, ****, P<0.0001. For **i-k**, data are representative of at least 3 experimental replicates and P values were calculated using a two-way ANOVA.

We found that this NEM derivatization indeed permitted the detection of mitochondrial cysteine (**Figure 5b and Extended Data Figure 4c**). Consistent with our previous findings^5^, cystine starvation promoted the rapid depletion of cellular cysteine (**Figure 5b and Extended Data Figure 4c**). The rate of exhaustion was similar between the cytosol and matrix, with a half-life of less than 30 minutes across compartments. In contrast, cellular GSH levels declined more gradually over the course of a 20h, with the matrix pool being marginally more stable than the cytosol following starvation (**Figure 5c and Extended Data Figure 4d**). We next evaluated the effect of CHAC1 knockdown on cysteine and GSH availability at 12h of cystine starvation. Whereas CHAC1 loss had no discernable effect on matrix cysteine availability, cytosolic cysteine was modestly spared in CHAC1 deficient cells (**Figures 5d and 5e**). We observed a more striking effect on cytosolic GSH levels, which were significantly elevated in the absence of CHAC1 (**Figure 5f**). This GSH sparing effect was also evident in the matrix, which exhibited similar trends towards increased GSH availability (**Figure 5g**). Together these data indicate that CHAC1 is a significant modulator of GSH availability under cystine starvation.

To interrogate if mitochondrial GSH is a determinant factor of Fe-S protein function under cysteine limitation, we restricted the matrix GSH pool by knocking out the predominant mitochondrial GSH transporter SLC25A39 (**Figure 5h**)^50, 51^. We found that like CHAC1, SLC25A39 knockout did not impair Fe-S protein function under cystine replete conditions (**Figures 5i-k**). However, the activities of ACO2 and the Fe-S dependent respiratory complexes were significantly diminished under cystine starvation in the absence of SLC25A39 (**Figures 5i-k**). Collectively these data indicate that CHAC1 catabolism of GSH has a significant influence on GSH availability in response to cystine starvation, and that mitochondrial GSH is critical for Fe-S protein function under these limiting conditions.

### Maintenance of Fe-S Clusters is Antagonistic to Survival Under Cystine Starvation

To determine if the GSH-dependent maintenance of Fe-S protein function plays a contributing role to ferroptosis in NSCLC we evaluated cell viability under cystine starvation in response to the disruption of Fe-S cluster homeostasis. Beyond synthesis, Fe-S clusters are regulated at the level of their oxidation status (**Figure 6a**)^52, 53^. We found that knockdown of components of the core Fe-S cluster biosynthetic complex diminished Fe-S protein function and prolonged survival of H1299 and PC9 cells starved of cystine (**Figure 6b and Extended Data Figures 5a-c**). A previous report found that loss of Fe-S cluster synthesis exacerbated erastin-induced ferroptosis in A549 cells due to an accumulation of iron^54^. Consistent with this report, we find that NFS1 knockdown provides no survival benefit under cystine starvation in A549 cells and that ISCU knockdown exacerbates ferroptosis in these cells (**Extended Data Figures 5a and 5d**). This suggests a cell line dependent response to the targeted disruption of the Fe-S cluster synthesis machinery under cystine starvation. We previously established that the mitochondrial NADPH producing nicotinamide nucleotide transhydrogenase (NNT) supports Fe-S protein function through the mitigation of cluster oxidation^37^. We observed that the decrease in respiratory chain function upon NNT knockdown was associated with an increase in survival under cystine deprivation (**Figure 6c and Extended Data Figures 5e-g**). Together these data suggest that the maintenance of Fe-S cluster homeostasis is detrimental to cystine starved NSCLC cells.

**Figure 6:**
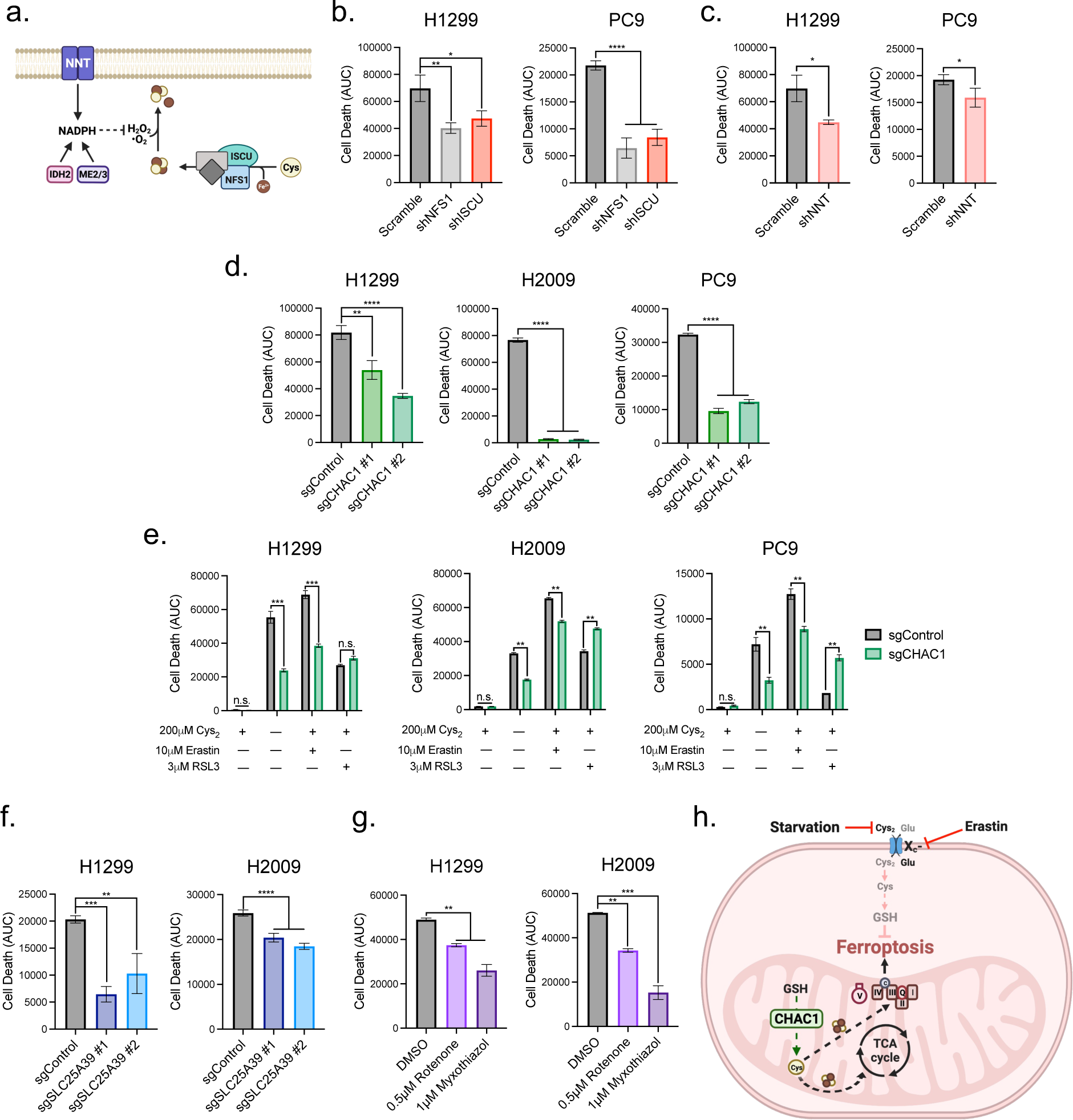
Mitochondrial Respiratory Function Antagonizes NSCLC Viability in the Absence of Extracellular Cys_2_. **a**, Schematic representation of the two major aspects of Fe-S cluster homeostasis: *de novo* synthesis and mitigation of cluster oxidation. **b** and **c**, Quantification of cell death over 48h of cystine starvation in H1299 and PC9 cells following the disruption of Fe-S cluster homeostasis through shRNA knockdown of **b**, NFS1 and ISCU, or **c**, NNT (n=3 per condition). **d**, Measurement of CHAC1-deficient H1299, H2009, and PC9 cell death under 48h of cystine starvation (n=3 per condition). **e**, Quantification of CHAC1-deficient H1299, H2009, and PC9 cell death over 48h of treatment with indicated inducers of ferroptosis (n=3 per condition). **f**, Determination of cell death in SLC25A39-deficient H1299 and H2009 cells starved of extracellular cystine for 48h (n=3 per condition). **g**, Quantification of cell death over 48h in H1299 and H2009 cells subject to ETC inhibition under cystine starvation (n=3 per condition). **h**, Summary schematic depicting that CHAC1 catabolism of GSH supports mitochondrial metabolic function when Cys is limiting to the detriment of NSCLC cell viability. Data represent mean values ± s.d; n.s., not significant, *, P<0.05, **, P<0.01, ***, P<0.001, ****, P<0.0001. For **b-g**, data are representative of at least 3 experimental replicates. For **b-d**, **f** and **g**, P values were calculated using a one-way ANOVA. For e, P values were calculated using a two-way ANOVA.

We next evaluated if the defect in Fe-S protein function observed with CHAC1 loss under cystine deprivation was also associated with a delay in ferroptosis. Indeed, CHAC1 knockout significantly prolonged survival under cysteine deprivation (**Figure 6d**). Further, this was associated with a decrease in plasma membrane lipid peroxidation (**Extended Data Figure 5h**). To extend this analysis, we determined if CHAC1 knockout influenced the response to other ferroptosis inducers. We found that CHAC1 knockout cells were similarly protected against erastin induction of ferroptosis, but not against the GPX4 inhibitor RSL3 (**Figure 6e**). This suggests that the effect of CHAC1 on ferroptosis induction is at least in part due to its effect on the mitochondria, as only cysteine restriction and not GPX4 inhibition-induced ferroptosis has an established mitochondrial component^13^. However, this is unlikely to be a result of CHAC1 sparing of matrix GSH, as SLC25A39 also prolonged survival under cysteine deprivation (**Figure 6f**). Finally, we employed inhibitors of complex I (rotenone) and complex III (myxothiazol) to confirm if the consequence of Fe-S protein function, sustained ETC flux, impacts the induction of ferroptosis. In support of this, we found that both rotenone and myxothiazol prolonged survival in the absence of cystine (**Figure 6g**). In totality, our findings indicate that NSCLC cells sustain their respiratory capacity in response to cysteine deprivation to the detriment of their viability. This is achieved through the maintenance of mitochondrial cysteine metabolism in the form of Fe-S cluster synthesis and depends on the catabolism of mitochondrial GSH by CHAC1 (**Figure 6h**).

## Discussion

Mitochondria are critical mediators of the cellular response to various stressors, integrating both intrinsic and extrinsic signals to modulate their function in support of cell resilience^55^. However, there exist instances in which mitochondria compete with the broader cellular milieu to selfishly benefit themselves at the level of the mitochondrial genome^56^. This is likely an ancient relic of endosymbiosis and has a significant impact on human disease^56–58^. Our findings indicate a selfish response to nutrient stress in NSCLC at the level of mitochondrial metabolism. Herein we describe a self-defeating maintenance of respiratory function through the persistence of mitochondrial cysteine metabolism in the absence of an extracellular source. This occurs through an apparent exploitation of the cellular stress response; where the activation of ATF4 in response to cystine starvation drives the expression of CHAC1^39, 41^.CHAC1 is recognized as a potential biomarker for ferroptosis^6, 59^ and has been implicated in execution of this cell death pathway^59^. However, the mechanism by which CHAC1 promotes ferroptosis has not been adequately defined. Our work demonstrates that CHAC1 catabolism of GSH is a significant factor in the depletion of GSH across cellular compartments. In the cytosol of NSCLC cells this is accompanied by an accumulation of ROS^5^, which facilitates peroxidation of plasma membrane lipids^8^. Intriguingly, we find that the depletion of matrix GSH is not associated with an accumulation of oxidative stress in NSCLC mitochondria. Instead, the regulated degradation of GSH by CHAC1 actually sustains mitochondrial function through the support of Fe-S cluster synthesis in the absence of a soluble pool. This finding agrees with a recent report uncovering a significant connection between mitochondrial GSH and Fe-S proteins at the organismal level^50^, and indicates an aspect of this regulation is at the level of cluster synthesis itself.

Yet, this persistence of mitochondrial function under cysteine limitation may be a unique characteristic of NSCLC compared to other malignancies. Though mitochondrial metabolism is implicated in ferroptosis, mitochondrial function is reported to progressively decline upon prolonged cysteine deprivation, marked by drastic changes in membrane potential, damage to the IMM, and fragmentation of the mitochondrial network in fibrosarcoma, breast and liver cancers^13, 20, 59^. Moreover, the systemic depletion of cysteine promoted severe mitochondrial morphological changes and loss of cristae in pancreatic ductal carcinoma^3^. The propensity for lung cancer cells to retain their mitochondrial function under this nutrient stress may be a factor of their tissue of origin. The lung experiences atmospheric oxygen tension upon inspiration, saturating the tissue with this vital nutrient^60^. However, oxygen is incredibly toxic in excess, and the mitochondria serve a critical function in coupling its detoxification to the generation of energy^61^. Alveolar type II cells, which are appreciated as a cell of origin for lung adenocarcinoma^62, 63^, are enriched with mitochondria to manage the highly oxygenated lung microenvironment^64^. This may contribute to the substantial mitochondrial metabolism exhibited by human lung tumors^23, 24^ and the necessity for mitochondrial function in lung tumorigenesis^37, 65–68^. Considering the importance of mitochondrial function in lung tumors, our data may reflect an inherent prioritization of mitochondrial function in response to stress in NSCLC.

Prioritization of mitochondrial function causes an apparent misalignment with the capacity to maintain cell viability under cysteine starvation, as our findings reinforce the connection between ETC activity and ferroptosis^13–16^. The connection between this critical aspect of mitochondrial metabolism and ferroptosis remains unresolved and is a significant focus of our ongoing work. Though we show that the maintenance of Fe-S protein function by CHAC1 activity in the mitochondria promotes ferroptosis, this does not preclude its effect on cytosolic GSH. We show that CHAC1 loss significantly spares cytosolic GSH under cystine starvation, indicating that GSH catabolism is a relevant factor in the capacity of cells to mitigate lipid peroxidation^10^. Although this is detrimental in the context of cysteine limitation, this explains why CHAC1 expression is restricted under basal conditions^39, 41, 69^. CHAC1 exhibits 20X the catalytic efficiency of CHAC2, which is constitutively expressed for the maintenance of GSH at millimolar levels within the cell^5, 70^. Therefore, CHAC1 likely serves a cytoprotective role when excess GSH is detrimental to the cell, such as conditions of reductive stress^71, 72^. In light of our findings, a characterization of CHAC1 function in response to various redox stressors is highly warranted.

## Methods

### Cell Culture

Human lung adenocarcinoma cell lines (A549, H322, H1299, H1792, H1975, H2009, PC9) were previously obtained^73^ from the Harmon Cancer Center Collection (University of Texas-Southwestern Medical Center) or commercially from the American Type Culture Collection (ATCC) and cultured in RPMI 1640 medium (Fisher Scientific, 11-875-119) supplemented with 5% FBS (Sigma-Aldrich, F0926) in the absence of antibiotics. Cell lines were maintained at 37°C in a humidified incubator containing 95% air and 5% CO_2_ and routinely tested for mycoplasma contamination with the MycoAlert Assay (Lonza, LT07-418). For experiments requiring manipulation of cystine availability, cells were cultured in RPMI 1640 medium without L-glutamine, L-cysteine, L-cystine, and L-methionine (Fisher Scientific, ICN1646454) supplemented with 5% dialyzed FBS (Sigma-Aldrich, F0392), L-glutamine (VWR, VWRL0131-0100), and L-methionine (Sigma-Aldrich, M5308). L-Cystine (Sigma-Aldrich, C6727) was also added back to the deficient media when required. Additional reagents used for the described experiments included: DMSO (VWR, 97063), FCCP (Sigma-Aldrich, C2920), ferrostatin-1 (Cayman Chemical, 17729), Z-VAD-FMK (Fisher Scientific, F7111), cyclosporin A (Fisher Scientific, NC9676992), deferoxamine (DFO; Sigma-Aldrich, D9533), erastin (Cayman Chemical, 17754), RSL3 (Cayman Chemical, 19288), rotenone (Sigma-Aldrich, R8875), myxothiazol (Neta Scientific, T5580).

### Viral Infection

For the generation of lentivirus, Lenti-X 293T cells (Clontech, 632180) were cultured in DMEM (Fisher Scientific, MT10013CV) supplemented with 5% FBS to 90% confluency. They were then co-transfected in the presence of polyethylenimine (Sigma-Aldrich, 408727) with 6μg of the plasmid of interest and 6μg of the packaging plasmids pCMV-VSV-G (Addgene, 8454) and pCMVdR8.2 dvpr (Addgene, 8455) in a 1:8 ratio. Cells were infected in the presence of 8μg/mL polybrene (Santa Cruz Biotechnology, sc-134220A) with lentivirus for 8h at an optimized dilution determined with the Lenti-X GoStix Plus protocol (Takara, 631280). CHAC1 and SLC25A39 knockout cells were generated using the pLenti-CRISPR-V2 plasmid (Addgene, 52961) encoding validated sgRNA sequences towards CHAC1 or SLC25A39^74^. Knockout cells were selected with 1μg/mL puromycin (Invivogen, ant-pr-1) for 5d prior to use for experimentation. Additionally, shRNA sequences targeting CHAC1^75^ or CHAC2 were cloned into a modified version of the doxycycline inducible LT3GEPIR vector^46^, where the puromycin resistance cassette was replaced with a zeocin resistance gene to generate cell lines with temporal control of CHAC1 depletion. These cells were selected in 100μg/mL zeocin (Invivogen, ant-zn-1) for 7d prior to use for experimentation. Finally, shRNA sequences targeting NFS1 (Addgene, 102963), ISCU (Addgene, 102972), NNT (Open Biosystems, TRCN0000028507), or a nontargeting control sequence (Scramble; Millipore Sigma, SHC002) in a pLKO.1 vector were used to disrupt the expression of proteins associated with Fe-S cluster homeostasis. Cells were selected in 1μg/mL puromycin for 3d and then used for experimental analysis 4d post-infection.

For the generation of 3XHA-EGFP-OMP25 (HA-Mito) retrovirus^44^, Phoenix-AMPHO 293T cells (ATCC, CRL-3213) were cultured in DMEM supplemented with 5% FBS to 90% confluency. They were then transfected with 6μg of the pMXs-3XHA-EGFP-OMP25 (Addgene, 83356) plasmid in the presence of polyethylenimine. Cells were infected in the presence of 8μg/mL polybrene with undiluted retrovirus for 24h, then overlaid with fresh retrovirus for an additional 24h. Retrovirally-infected cells were then selected with 10μg/mL blasticidin (Invivogen, ant-bl-1) for 5d prior to use for experimentation.

### Seahorse Analyses of Mitochondrial Function

Analyses of mitochondrial metabolic function were performed with a Seahorse XFe96 Analyzer (Agilent). General mitochondrial function was assessed based on the Seahorse XF Cell Mito Stress Test protocol (Agilent Technologies, Santa Clara, CA, USA). Basal oxygen consumption rate (OCR) was determined by subtracting rotenone and antimycin A (Sigma-Aldrich, A8674) insensitive oxygen consumption from baseline measurements. Coupling efficiency was determined by calculating the proportion of rotenone/antimycin A sensitive oxygen consumption that is also sensitive to oligomycin (Sigma-Aldrich, 75351) treatment. ATP-linked respiration was determined by calculating the fraction of baseline oxygen consumption that is sensitive to ATP synthase inhibition. Finally, maximal respiratory capacity was determined by subtracting the residual oxygen consumption following rotenone/antimycin A treatment from the oxygen consumption stimulated by mitochondrial uncoupling. Fe-S cluster dependent ETC complex function was assessed according to a previously established protocol^36^. Briefly, 40,000 cells were seeded overnight on quadruplicate wells of an XFe96 microplate. Cells were then washed twice with mitochondrial assay solution (220mM mannitol [Sigma-Aldrich, M4125], 70mM sucrose [Sigma-Aldrich, S7903], 10mM KH_2_PO_4_ [VWR, 470302], 2mM HEPES [Fisher Scientific, BP310], and 1mM EGTA [VWR, 97062]) and then overlaid with 175μL of mitochondrial assay solution supplemented with 10mM sodium pyruvate (Sigma-Aldrich, P5280), 1mM malate (Sigma-Aldrich, M0875), 4mM ADP (Sigma-Aldrich, A5285), and Seahorse Plasma Membrane Permeabilizer (Agilent, 102504-100). Cells were then sequentially subjected to 1μM rotenone to inhibit complex I, 10mM succinate (Sigma-Aldrich, S3674) to stimulate complex II, and finally 1μM antimycin A to inhibit complex III. For assays requiring a period of cystine starvation, 20,000 cells were seeded overnight and then overlaid with cystine replete or deficient RPMI for 20 hours prior to starting the assay. Following each assay, cells in each well were lysed to determine protein abundance for normalization.

### Cell Viability

Cells were seeded overnight onto black walled 96-well plates at a density of 7,500-12,500 cells/well in a total volume of 100μL. The cells were then washed twice with PBS and then overlaid with cystine-free RPMI supplemented with 20nM Sytox Green (Fisher Scientific, S7020) and the indicated treatment. Cells were placed in an IncuCyte S3 live-cell analysis system (Essen BioScience, Ann Arbor MI, USA) or a CellCyte X Live Cell Analyzer (Cytena, Freiburg im Breisgau, Germany) housed in a humidified incubator containing 95% air and 5% CO_2_ at 37°C. A series of images were acquired every 2-4h with a 10X objective lens in phase contrast and green fluorescence (Ex/Em: 460/524nm with an acquisition time of 200-400ms). Data was processed using IncuCyte S3 2020A (Essen BioScience, Ann Arbor MI, USA) or CellCyte Studio (Cytena, Freiburg im Breisgau, Germany) software. Cell death was calculated as the number of Sytox Green positive cells/mm^2^ and normalized to cell confluence (dead cells/mm^2^/% confluence). Cell death data were represented as area under the curve (AUC), which represents the sum of dead cells over a 48h period.

### Fluorescence-Based Analyses of ROS

Fluorescent probes assessing ΔΨ_m_, mitochondrial density, cellular ROS, mitochondrial •O_2_^-^, H_2_O_2_, and lipid peroxidation were measured by flow cytometry. For each treatment condition, 10^5^ cells were seeded overnight on triplicate wells of a 6-well plate. Cells were then washed twice with PBS and overlaid with cystine replete or deficient RPMI for 20 hours. ΔΨ_m_ was determined using tetramethylrhodamine (TMRE; Fisher Scientific, T669) as previously described^76^. At 19.5h of treatment, cells were incubated in 250nM TMRE suspended in the appropriate treatment media for 30min at 37°C and then collected for analysis. Mitochondrial density was determined using MitoTracker Green FM (ThermoFisher Scientific, M7514) according to the manufacturer’s protocol. Following treatment, cells were incubated in 200nM MitoTracker Green FM in PBS for 15min at 37°C and then collected for analysis. Cellular ROS levels were assayed using CellROX Green (ThermoFisher Scientific, C10444) according to the manufacturer’s protocol. Briefly, at 19.5h of treatment, 4μL of 2.5mM CellROX Green was applied to each well for a working concentration of 5μM. Cells were incubated at 37°C for the remaining 30min of treatment and then collected for analysis. Mitochondrial •O_2_^-^ was determined with MitoSOX Red (ThermoFisher Scientific, M36008) based on a previously described protocol^77^. Following treatment, cells were incubated in 5μM of MitoSOX Red in PBS for 20min at 37°C and then collected for analysis. Mitochondrial H_2_O_2_ was assessed with MitoPY1 (Fisher Scientific, 442810) according to an established protocol^78^. Following treatment, cells were incubated in 10μM of MitoPY1 in PBS for 20min at 37°C and then collected for analysis. Mitochondrial lipid peroxidation was determined using the ratiometric fluorescent probe MitoPerOX (Cayman Chemical, 18798) according to the manufacturer’s protocol. At 19.5h of treatment, cells were incubated in 10μM MitoPerOX suspended in the appropriate treatment media for 30min at 37°C and then collected for analysis. For all analyses, cells were washed twice with ice cold PBS following incubation with the respective probe and collected into 500μL of ice cold PBS for analysis with a BD Accuri C6 Plus Flow Cytometer (BD Biosciences, Franklin Lakes, NJ, USA). For TMRE, and MitoSOX Red, a phycoerythrin (PE) optical filter was used for fluorescence detection. For MitoTracker Green, CellROX Green, and MitoPY1, a fluorescein isothiocyanate (FITC) optical filter was used for fluorescence detection. MitoPerOX fluorescence was detected with both the PE and FITC filters to calculate the ratio of oxidized (green) and unaltered (red) membrane lipids. For each analysis, the mean fluorescence intensity of 10,000 discrete events was calculated. Whole cell lipid peroxidation was determined using the Image-IT Lipid Peroxidation Kit (ThermoFisher Scientific, C10445) according to the manufacturer’s protocol. For each treatment condition, 15,000 cells were seeded overnight onto triplicate wells of a black walled 96-well plate. Cells were then washed with PBS and treated with cystine replete or deficient RPMI for 20h. At 19.5h of treatment, 2μL of 10mM BODIPY-C11 was applied to each well for a working concentration of 10μM. Cells were incubated at 37°C for the remaining 30min of treatment. Cells were washed twice with PBS and then placed in a fluorescence-compatible plate reader and fluorescence measured (reduced Ex:475nm/Em:580-640nm, oxidized Ex:475nm/Em:500-550nm).

### Mitochondrial GSH:GSSG Ratio

The oxidation state of the mitochondrial GSH pool was assessed using a previously developed genetically encoded fluorescent and ratiometric biosensor targeted to the mitochondria^32^ (Mito-Grx-roGFP2). The Mito-Grx1-roGFP2 construct (Addgene, 64977) was cloned into the pLenti-CMV-Puro vector (Addgene, 17448) and lentivirus generated to produce stable cell lines expressing the biosensor as described. For each treatment condition, 15,000 cells were seeded overnight onto triplicate wells of a black walled 96-well plate. Cells were then washed with PBS and treated with cystine replete or deficient RPMI for 16 hours. Cells were then placed in a fluorescence-compatible plate reader (Promega, Madison, WI, USA) and fluorescence measured (GSH-bound Ex:475nm/Em:500-550nm, GSSG-bound Ex:405nm/Em:500-550nm).

### Immunoblotting

Following the indicated treatment, cell lysates were generated in ice cold RIPA lysis buffer (20mM Tris-HCl (VWR, 97061-258), pH 7.5, 150mM NaCl (Fisher Scientific, S271), 1mM EGTA, 1mM EDTA [Sigma-Aldrich, E5134], 1% sodium deoxycholate [Sigma-Aldrich, D6750], 1% NP-40 [Sigma-Aldrich, 74385]) supplemented with protease inhibitors (Fisher Scientific, PIA32955). Protein concentrations were then determined by DC Protein Assay (Bio-Rad, 5000112) and 10-30μg samples were prepared with a 6x reducing sample buffer containing β-mercaptoethanol (VWR, M131). Proteins were resolved by SDS-PAGE using NuPAGE 4-12% Bis-Tris precast gels (Fisher Scientfic, WG1402BOX) and then transferred to 0.45μM nitrocellulose membranes (VWR, 10120-006). Membranes were then blocked with 5% nonfat dairy milk in Tris-buffered saline containing 0.1% Tween 20 (VWR, 0777), and incubated overnight with the following primary antibodies: ACC (1:1000; Cell Signaling Technologies, 3662S, RRID:AB_2219400), ATF4 (1:1000; Cell Signaling Technologies, 11815, RRID:AB_2616025), CHAC1 (1:1000; Proteintech, 15207-1-AP, RRID:AB_2878118), CHAC2 (1:1000; GeneTex, GTX128819, RRID:AB_2885821), DLST (1:1000; Cell Signaling Technologies, 11954S, RRID:AB_2732907), ERp57 (1:1000; Cell Signaling Technologies, 2881, RRID:AB_2160840), HSP60 (1:1000; Cell Signaling Technologies, 4870S, RRID:AB_2295614), HSP90 (1:1000; Cell Signaling Technologies, 4874S, RRID:AB_2121214), ISCU (1:1000; Santa Cruz Biotechnology, sc-373694, RRID:AB_10918261), Lipoic Acid (1:1000; Millipore Sigma, 437695-100UL, RRID:AB_10683357), MCU (1:1000; Cell Signaling Technologies, 14997S, RRID:AB_2721812), NFS1 (1:1000; Santa Cruz Biotechnology, sc-365308, RRID:AB_10843245), NNT (1:1000; Abcam, ab110352, RRID:AB_10887748), PDH-E2 (1:1000; Abcam, ab126224, RRID:AB_11129511), PRDX3 (1:1000; Abcam, ab73349, RRID:AB_10843245), NNT (1:1000; Abcam, ab110352, RRID:AB_10887748), PDH-E2 (1:1000; Abcam, ab126224, RRID:AB_11129511), PRDX3 (1:1000; Abcam, ab73349, RRID:AB_1860862), SHMT2 (1:1000; Cell Signaling Technologies, 12762S, RRID:AB_2798018), SLC25A39 (1:1000; Proteintech, 14963-1-AP, RRID:AB_2878095), TOM20 (1:1000; Santa Cruz Biotechnology, sc-17764, RRID:AB_628381), α-Tubulin (1:1000; Abcam, ab7291, RRID:AB_2241126). HRP-conjugated anti-mouse (1:10,000; Jackson ImmunoResearch, 115-005-003, RRID:AB_2338447) and anti-rabbit (1:10,000; Jackson ImmunoResearch, 111-005-003, RRID:AB_2337913) secondary antibodies and enhanced chemiluminescence were then used for all immunoblotting.

For the determination of the PRDX3 oxidation state, a previously established redox immunoblotting protocol was employed^79^. Briefly, following the indicated treatment, cells were washed twice with PBS and overlaid with 200 μL of alkylation buffer (40 mM HEPES, 50 mM NaCl, 1 mM EGTA, protease inhibitors) supplemented with 25 mM N-ethylmaleimide (NEM; Fisher Scientific, AA4052606). Cells were incubated for 10 minutes at room temperature and then 20 μL of 10% CHAPS detergent (Sigma-Aldrich, C3023) was added and cells incubated at room temperature for an additional 10 minutes to lyse cells. Lysates were then cleared by centrifugation at 4 °C for 15 minutes at 17,000 *g*. Supernatants were isolated to quantify protein and 5-10 μg protein samples were mixed with a 4X non-reducing buffer prior to separation by SDS-PAGE as described.

### ACO2 Activity

Mitochondrial aconitase (ACO2) activity was determined as we have previously described^37^. Briefly, cystine fed or starved cells were collected and resuspended in 200μL of 50mM Tris-HCl and 150mM NaCl, pH 7.4. Cells were then homogenized by dounce homogenizer (15 strokes) and the homogenate spun down at 4°C for 10min at 10,000 *g*. Organellar pellet was then washed twice and resuspended in 100μL of 1% Triton X-100 (VWR, 0694) in 50mM Tris-HCl and 150mM NaCl, pH 7.4 to lyse the mitochondrial membrane. Lysates were then spun down at 4°C for 15 minutes at 17,000 *g*. Protein concentrations were then determined by DC Protein Assay, and 175μL of 100-500 μg/mL protein were generated in 50mM Tris-HCl, pH 7.4 and incubated at 37°C for 15 minutes. 50μL of this protein solution was transferred to triplicate wells of a black walled 96-well plate containing 50μL of 50mM Tris-HCl, pH 7.4. Next, 50μL of a 4mM NADP^+^ (Neta Scientific, SPCM-N1131-OM) and 20 U/mL IDH1 (Sigma-Aldrich, I1877) solution and 50μL of 10mM sodium citrate were sequentially added to initiate the assay. The plate was transferred to a fluorescence-compatible plate reader maintained at 37°C to measure the change in NADPH autofluorescence every minute over a period of 45 minutes. The accumulation of NADPH autofluorescence over time is an indicator of aconitase activity, where ACO2 present in the sample converts the supplied citrate to isocitrate, which is subsequently metabolized by the supplied IDH1 in a reaction that yields NADPH.

### Mitochondrial Immunoprecipitation

An established protocol for the rapid isolation of mitochondria from HA-Mito cells^44^ was modified as follows to preserve thiol containing metabolites and permit their detection by LC-MS. Prior to mitochondrial isolation, 50-200μL of Pierce Anti-HA magnetic beads (VWR, PI88837) were washed in KPBS (136mM KCl [Sigma-Aldrich, P5405], 10mM KH_2_PO_4_, pH 7.4 in HPLC grade H_2_O [Fisher Chemical, W5-1]) for each sample. Samples were processed one at a time in a 4°C cold room to best preserve the metabolic state of each sample. Cells were washed twice with ice cold KPBS and then incubated in KPBS supplemented with 25mM NEM and 10mM ammonium formate (Fisher Scientific, 501454965) for 1min. Cells were then washed twice more and collected into ice cold KPBS. Cells were pelleted at 1,000 *g* for 2min and then resuspended in 200μL of KPBS. Cells were homogenized by Dounce homogenizer (25 strokes), and the homogenate spun down at 1,000 *g* for 2min. The supernatant was then applied to the pre-washed magnetic beads and incubated on an end-over-end rotator for 3.5min. Beads were then isolated from solution with a DynaMag-2 magnetic stand (ThermoFisher Scientific, 12321D). The unbound solution was kept as a mitochondrial-free cytosolic fraction. Mitochondria-bound beads were washed twice with KPBS and then processed for either immunoblotting or metabolite detection. For the purposes of immunoblotting, isolated beads were reconstituted in 50μL of lysis buffer (50mM Tris-HCl, pH 7.4, 150mM NaCl, 1mM EDTA, 1% Triton X-100, protease inhibitors), incubated on ice for 10min and lysates cleared by centrifugation at 17,000 *g* for 10 minutes. Protein was quantified and samples prepared for SDS-PAGE. For metabolite detection, beads were reconstituted in 50μL of 80% methanol containing 20 μM of [^13^C_3_, ^15^N]-NEM-Cysteine (Cambridge Isotope Laboratories, CNLM-38710H-0.25) and 40 μM of [^13^C_2_, ^15^N]-NEM-GSH (Sigma-Aldrich, 683620), incubated on ice for 5 minutes, and extracts cleared by centrifugation at 17,000 *g* for 10 minutes. NEM-derivatized standards were prepared as previously described^5^. Alternatively, the mitochondrial-free cytosolic fractions were extracted in 800μL of the defined extraction solution, incubated on ice for 5 minutes, and cleared by centrifugation at 17,000 *g* for 10 minutes.

### LC-MS-Based Metabolite Detection

For determination of the relative Cys and GSH pool sizes across compartments, previously established LC-MS conditions were employed^5, 22^ For chromatographic metabolite separation, a Vanquish UPLC system was coupled to a Q Exactive HF (QE-HF) mass spectrometer equipped with HESI (Thermo Fisher Scientific, Waltham, MA, USA). Samples were run on an Atlantis Premier BEH Z-HILIC VanGuard FIT column, 2.5µm, 2.1mm × 150mm (Waters, Milford, MA, USA). The mobile phase A was 10mM (NH_4_)_2_CO_3_ and 0.05% NH_4_OH in H_2_O, while mobile phase B was 100% ACN. The column chamber temperature was set to 30°C. The mobile phase condition was set according to the following gradient: 0-13min: 80% to 20% of mobile phase B, 13-15min: 20% of mobile phase B. The ESI ionization mode was negative, and the MS scan range (m/z) was set to 65-975. The mass resolution was 120,000 and the AGC target was 3 × 10^6^. The capillary voltage and capillary temperature were set to 3.5 kV and 320°C, respectively. 5μL of sample was loaded. The LC-MS metabolite peaks were manually identified and integrated by El-Maven (Version 0.3.1) by matching with a previously established in-house library^5^.

### Statistical Analysis

All data were analyzed for statistical significance with GraphPad Prism 9.4.1 software; P values <0.05 were deemed significant (n.s., not significant, *, P<0.05, **, P<0.01, ***, P<0.001, ****, P<0.0001). Comparisons of two groups were performed with a two-tailed unpaired Student’s *t*-test. For comparisons of a single variable in 3 or more groups a one-way ANOVA with a Dunnet’s multiple comparisons test was performed. For comparisons of two variables a two-way ANOVA with a Sidak’s multiple comparisons test was performed.

## Data Availability

All data supporting the findings of this study are available from the corresponding authors on request.

## Acknowledgements

We would like to thank members of the DeNicola and Ana Gomes laboratories for their very helpful discussions. Further, we thank the laboratories of John Cleveland, Elsa Flores, Bob Gillies, Ana Gomes, and Vince Luca for technical support in carrying out this study. This work was supported by grants from the NIH/NCI (R37CA230042 and P01CA250984) to G.M.D. This work was also supported by the Proteomics/Metabolomics Core, which is funded in part by Moffitt’s Cancer Center Support Grant (NCI, P30-CA076292). All schematics were created with BioRender.com.

## Contributions

N.P.W. conceived the study, designed and performed the experiments, analyzed the data, and wrote the manuscript; S.J.Y. performed LC-MS analysis of thiol metabolites; T.F. generated CHAC1-knockout lines and performed immunoblotting; A.S. generated SLC25A39-knockout lines and performed immunoblotting, J.M. characterized CHAC1 localization and expression in response to Cys_2_ starvation; G.M.D. conceived the study, acquired funding, contributed to the experimental design, supervised the project, and wrote the manuscript.

---

**Extended Data Figure 1:**
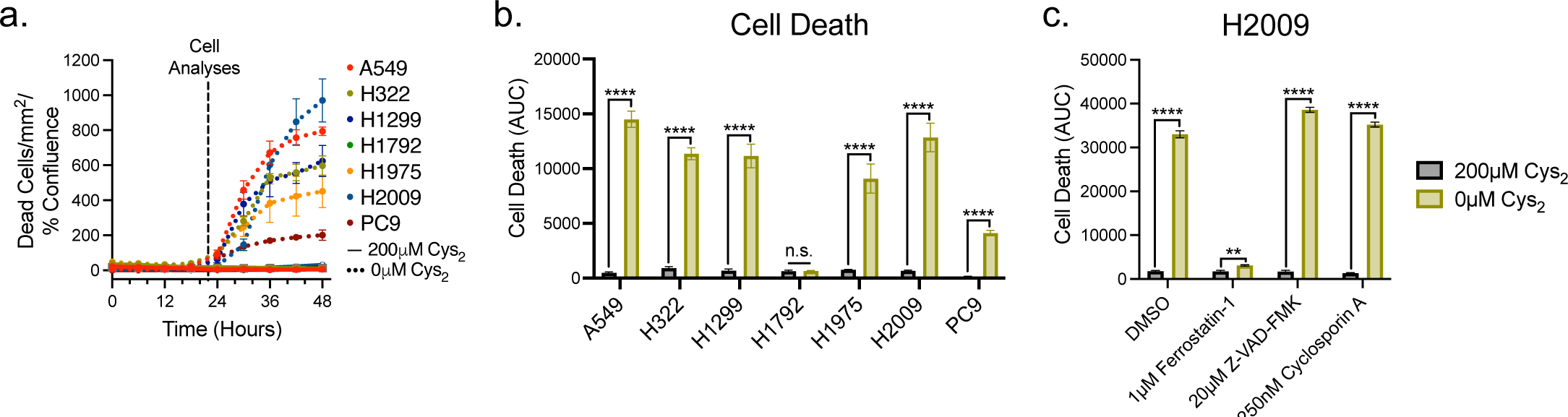
NSCLC Viability is Susceptible to Cys_2_ Starvation. **a**, Plot of the accumulation of dead NSCLC cells over time in cystine replete (solid) or starved (dashed) conditions (n=3 per condition). **b**, Measurement of NSCLC cell death over 48 hours in the presence or absence of cystine. Data represented as area under the curve (AUC) of the plots depicted in **a** (n=3 per condition). **c**, Measurement of cell death over 48 hours in H2009 cells fed/starved of cystine and treated with the indicated inhibitors of ferroptosis (Ferrostatin-1), apoptosis (Z-VAD-FMK), or necroptosis (Cyclosporin A) (n=3 per condition). Data represent mean values ± s.d; n.s., not significant, **, P<0.01, ****, P<0.0001. For **b**, P values were calculated using a two-tailed unpaired Student’s *t*-test. For **c**, P values were calculated using a two-way ANOVA. All data are representative of at least 3 experimental replicates.

**Extended Data Figure 2:**
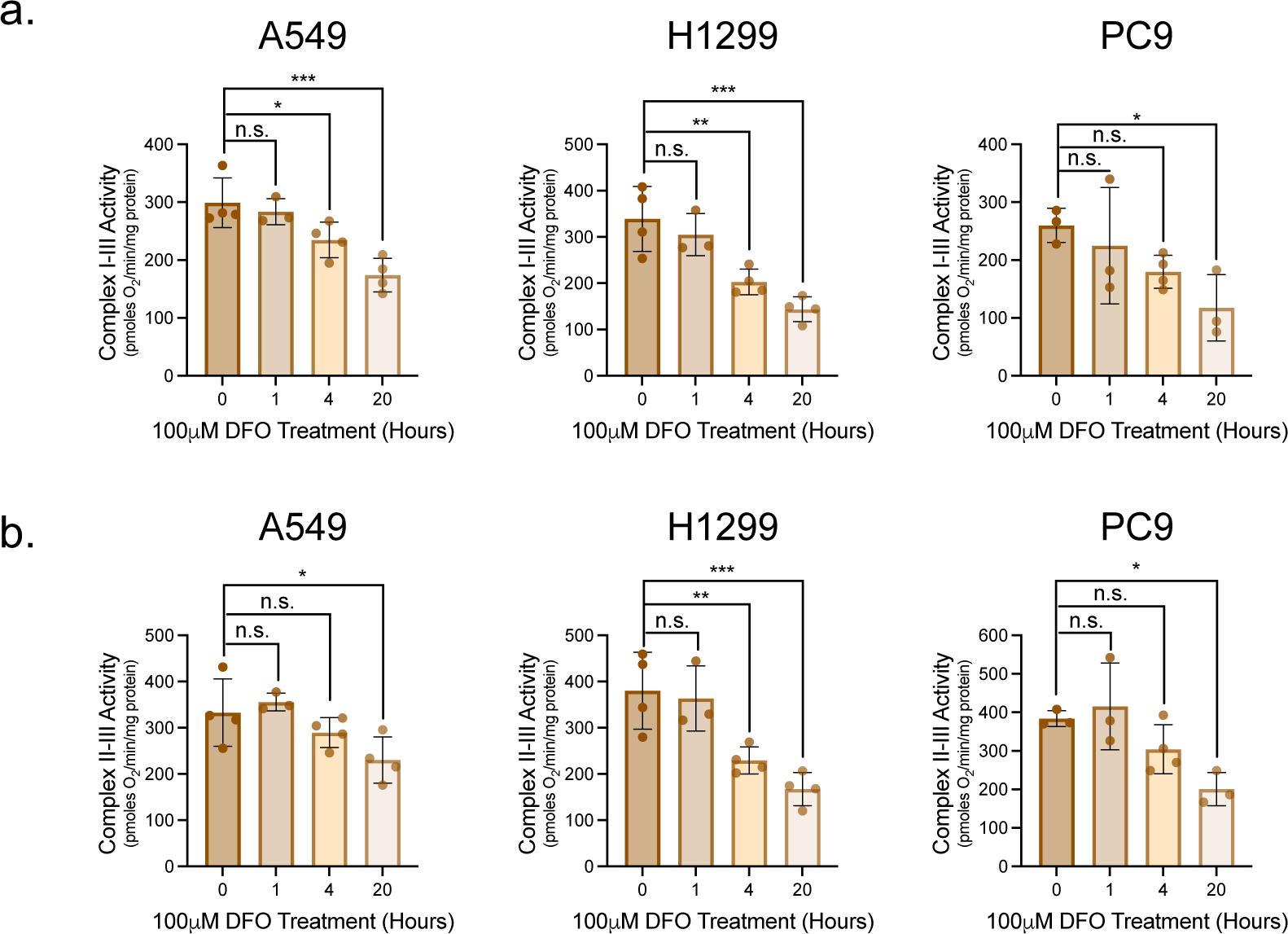
Iron Chelation Promotes a Gradual Loss of Fe-S Protein Function. **a**, Supercomplex I-III activity in permeabilized A549, H1299, and PC9 cells stimulated with 10mM pyruvate and 1mM malate following treatment with DFO for the indicated time (n :=3 per condition). **b**, Supercomplex II-III activity in permeabilized A549, H1299, and PC9 cells subject to rotenone inhibition of complex I and stimulated with 10mM succinate following iron chelation for the indicated time (n :=3 per condition). Data represent mean values ± s.d; n.s., not significant, *, P<0.05, **, P<0.01, ***, P<0.001. For **a** and **b**, P values were calculated using a one-way ANOVA. All data are representative of at least 3 experimental replicates.

**Extended Data Figure 3:**
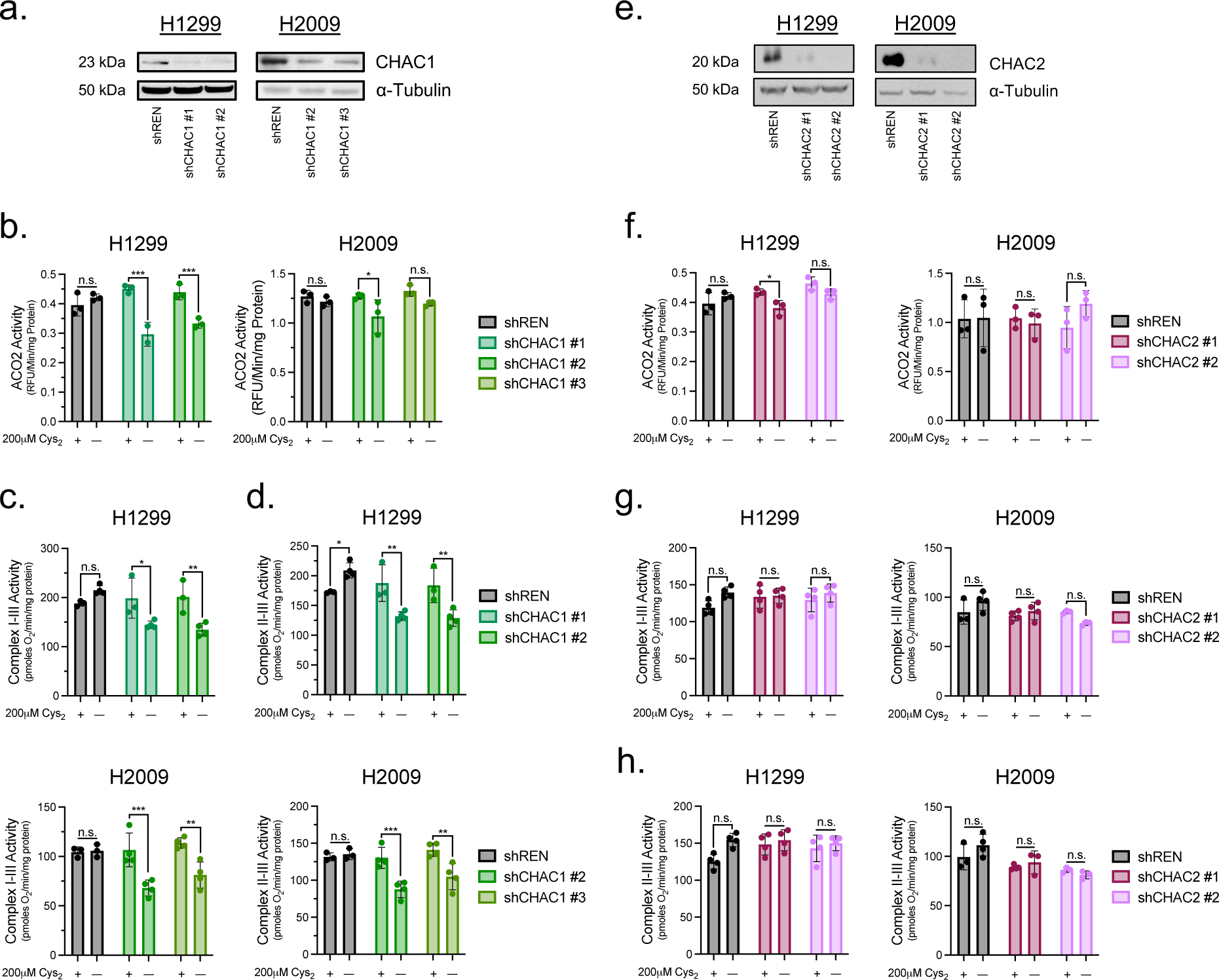
CHAC2 Loss Does Not Influence Fe-S Protein Function. **a**, immunoblot analysis of CHAC1 and α-Tubulin expression in H1299 and H2009 cells subject to 5-days of 0.2μg/mL doxycycline treatment to induce shRNA-mediated knockdown of CHAC1. **b**, ACO2 activities in H1299 and H2009 CHAC1-knockdown cells subject to either cystine replete or starved conditions (n=3 per condition). **c** and **d**, ETC supercomplex activities in permeabilized CHAC1-knockdown cells stimulated with **c**, 10mM pyruvate and 1mM malate or **d**, 10mM succinate following culture in the presence of absence of cystine (n :=3 per condition). **e**, immunoblot analysis of CHAC2 and α-Tubulin expression in H1299 and H2009 cells subject to 5-days of 0.2μg/mL doxycycline treatment to induce shRNA-mediated knockdown of CHAC2. **f**, ACO2 activities in H1299 and H2009 CHAC2-knockdown cells cultured in the presence or absence of cystine (n=3 per condition). **g** and **h**, ETC supercomplex activities in permeabilized CHAC2-knockdown cells stimulated with their associated substrates following culture in cystine replete or starved conditions (n :=3 per condition). For **b-d**, **f-h**, cells were treated with 0.2μg/mL doxycycline for 5 days prior to analysis and cultured in the indicated [cystine] for the final 20 hours. Data represent mean values ± s.d; n.s., not significant, *, P<0.05, **, P<0.01, ***, P<0.001, ****, P<0.0001. For **b-d, f-h**, data are representative of at least 3 experimental replicates and P values were calculated using a two-way ANOVA.

**Extended Data Figure 4:**
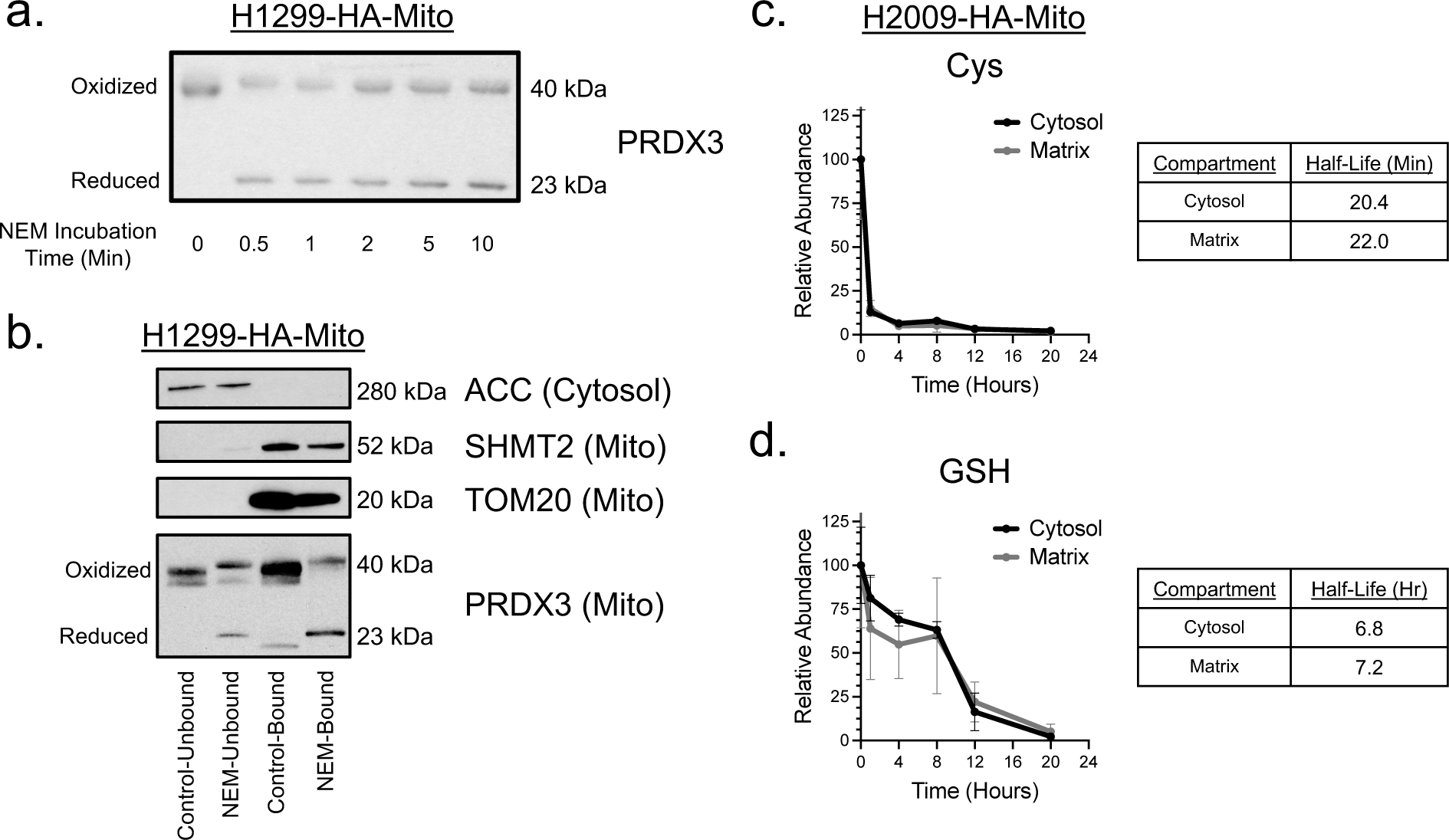
Cys_2_ Starvation Promotes a Rapid Depletion of Intracellular Cys. **a**, Redox immunoblot determination of reduced PRDX3 stabilization in H1299-HA-Mito cells treated with 25mM NEM for up to 10 minutes prior to mitochondrial immunoprecipitation. **b**, Immunoblot analysis of ACC, PRDX3, SHMT2, TOM20 expression in H1299-HA-Mito cells following mitochondrial immunoprecipitation ± a 1 minute incubation with 25mM NEM prior to mitochondrial isolation. **c**, Determination of the half-life of cytosolic and matrix Cys in H2009-HA-Mito cells cultured in the absence of extracellular Cys for 20h (n=3 biological replicates per time point). **d**, Determination of the half-life of cytosolic and matrix GSH in H2009-HA-Mito cells cultured in the absence of extracellular Cys for 20h (n=3 biological replicates per time point). Data represent mean values ± s.d.

**Extended Data Figure 5:**
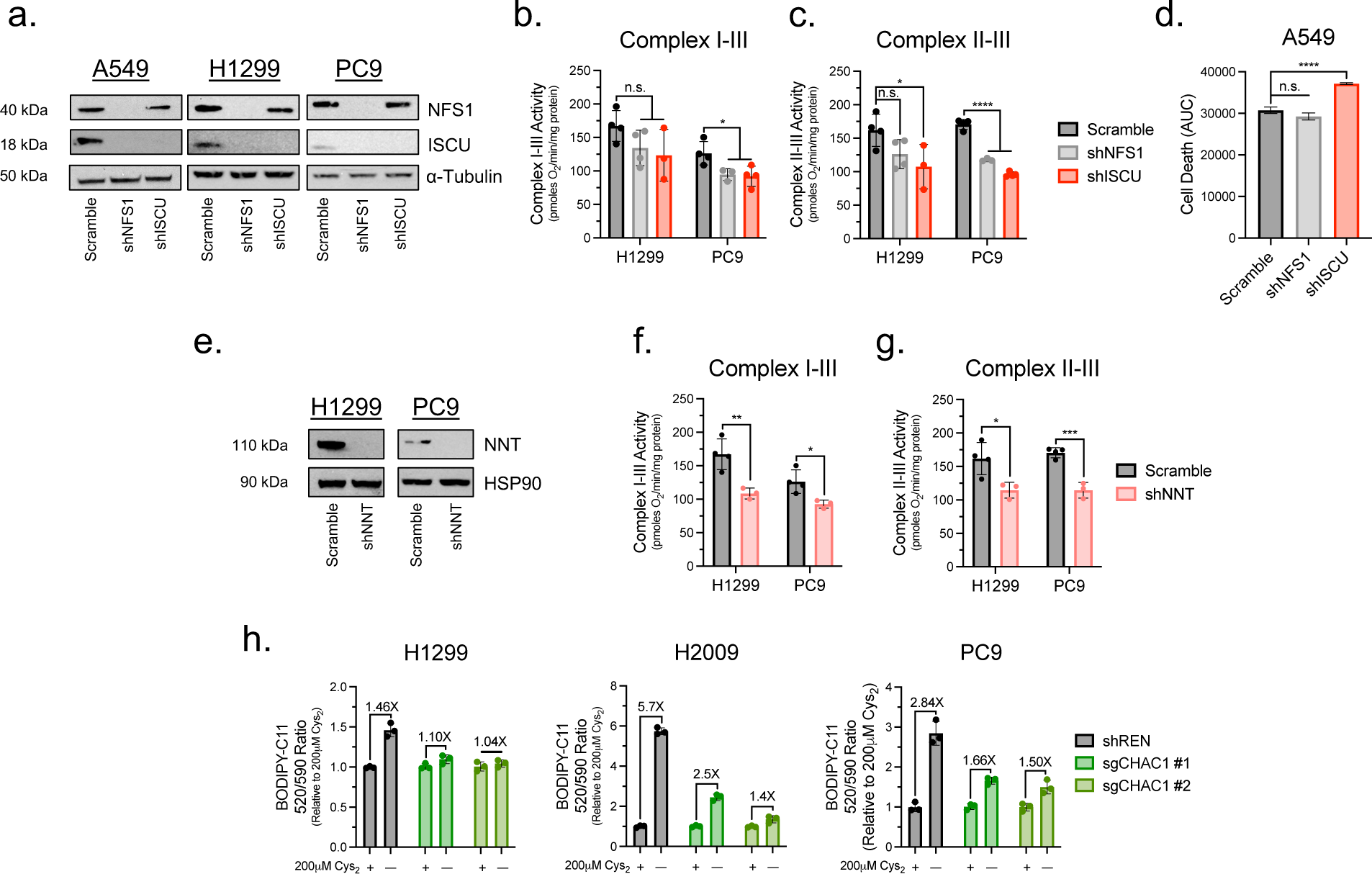
Fe-S Protein Function is Diminished Upon Disruption of Fe-S Cluster Homeostasis. **a**, Immunoblot analysis of NFS1, ISCU, and α-Tubulin expression in A549, H1299, and PC9 cells 4d post-infection with scramble, shNFS1, or shISCU lentivirus. **b** and **c**, ETC supercomplex activities in permeabilized H1299 and PC9 cells stimulated with **b**, 10mM pyruvate and 1mM malate or **c**, 10mM succinate 4d post-infection with scramble, shNFS1, or shISCU lentivirus (n :=3 per condition). **d**, Quantification of cell death over 48h of cystine starvation in A549 cells following the disruption of Fe-S cluster synthesis through knockdown of NFS1 and ISCU (n=3 per condition). **e**, Immunoblot analysis of NNT and HSP90 expression in H1299, and PC9 cells 4d post-infection with scramble or shNNT lentivirus. **f** and **g**, ETC supercomplex activities in permeabilized H1299 and PC9 cells stimulated with their associated substrates 4d post-infection with scramble or shNNT lentivirus (n :=3 per condition). **h**, Relative BODIPY-C11 fluorescence ratio of CHAC1-deficient H1299, H2009, and PC9 cells cultured in the presence or absence of cystine for :=20h as an indicator of the extent of cellular lipid peroxidation (n=3 per condition). Data represent mean values ± s.d; n.s., not significant, *, P<0.05, **, P<0.01, ***, P<0.001, ****, P<0.0001. For **b-d,** P values were calculated using a one-way ANOVA. For **f** and **g**, P values were calculated using a two-tailed unpaired Student’s *t*-test. Data are representative of at least 3 experimental replicates.

